# Genetic analysis of *Caenorhabditis elegans pry-1/Axin* suppressors identifies genes involved in reproductive structure development, stress response, and aging

**DOI:** 10.1101/2021.05.19.444814

**Authors:** Avijit Mallick, Nikita Jhaveri, Jihae Jeon, Yvonne Chang, Krupali Shah, Hannah Hosein, Bhagwati P. Gupta

**Affiliations:** Department of Biology, McMaster University, 1280 Main Street West, Hamilton, ON, L8S4K1

## Abstract

The Axin family of scaffolding proteins regulates a wide array of developmental and post-developmental processes in eukaryotes. Studies in the nematode, *Caenorhabditis elegans*, have shown that the Axin homolog, PRY-1, plays essential roles in multiple tissues. To understand the genetic network of *pry-1*, we focused on a set of genes that are differentially expressed in the *pry-1*-mutant transcriptome and are linked to reproductive structure development. Eight of the genes (*ard-1, rpn-7, cpz-1, his-7, cdk-1, rnr-1, clsp-1*, and *spp-1*), when knocked down by RNA interference, efficiently suppressed the plate-level multivulva phenotype of *pry-1* mutants. In every case, other than *clsp-1* and *spp-1*, the ectopic vulval precursor cell (VPC) induction was also inhibited. The suppressor genes are members of known gene families in eukaryotes and perform essential functions. Our genetic interaction experiments revealed that except for *clsp-1*, the genes participate in one or more *pry-1*-mediated biological events. While four of them (*cpz-1, his-7, cdk-1*, and *rnr-1*) function in VPC induction, stress response, and aging, the other three (*spp-1, ard-1*, and *rpn-7*) are specific to one or more of these processes. Further analysis of the genes involved in aging showed that *his-7, cdk-1*, and *rnr-1* also interacted with *daf-16/FOXO*. The results of genetic epistasis experiments suggested that *his-7* functions upstream of *daf-16*, whereas *cdk-1* and *rnr-1* act downstream of the *pry-1-daf-16* pathway. Altogether, these findings demonstrate the important role of *pry-1* suppressors in *C. elegans*. Given that all of the genes described in this study are conserved, future investigations of their interactions with Axin and their functional specificity promises to uncover the genetic network of Axin under normal and disease states.

## INTRODUCTION

Much of genetic research is aimed at linking genes to phenotypes and understanding how changes in gene function affect biological processes. Studies in animal models have demonstrated that genes exert their effects through interactions with other genes to form gene networks. Disruptions of the activity of network components can result in disease development. Therefore, a comprehensive understanding of gene interaction networks is crucial for the discovery of effective treatments. Our group is currently investigating the genetic network of an Axin family member in the nematode, *Caenorhabditis elegans*. Axins are scaffolding proteins that play crucial roles in regulating conserved processes in metazoans; they integrate inputs from multiple interactors to coordinate downstream cellular signaling events. Moreover, Axin mutations have been implicated in multiple abnormalities and diseases (Mallick *et al*. 2019b). Therefore, elucidating the Axin signaling cascade can enhance our understanding of disease progression and the pathway could be an attractive therapeutic target.

Work from our lab and others have shown that the *C. elegans* Axin homolog PRY-1 is involved in multiple developmental and post-developmental processes, including embryogenesis, neuronal differentiation, vulva formation, lipid metabolism, and lifespan maintenance (Mallick *et al*. 2019b, 2020; Mallick and Gupta 2020). Initial studies on *pry-1* showed that the gene product acts as a negative regulator of canonical WNT signaling (Korswagen *et al*. 2002). PRY-1 forms a destruction complex in the absence of a WNT ligand, leading to the inhibition of the WNT effector, the β-catenin homolog BAR-1. Mutations in *pry-1* mimic activated WNT signaling and cause the translocation of BAR-1 to the nucleus, thereby promoting the expression of target genes (Gleason *et al*. 2002).

During vulval development, *pry-1* restricts the number of induced vulval precursor cells (VPCs). In a normal worm, three (P5.p, P6.p, and P7.p) of the eight Pn.p cells (n = 3-8), termed as the VPCs, participate in the formation of the vulva (Sulston and Horvitz 1977; Sternberg 2005). The remaining uninduced VPCs adopt non-vulval fates and fuse with the surrounding hypodermal syncytium (Sternberg and Horvitz 1989). In the absence of *pry-1*, non-vulval cells are inappropriately induced to adopt specific fates, resulting in multiple ectopic ventral protrusions, a phenotype termed multivulva (Muv) (Gleason *et al*. 2002; Seetharaman *et al*. 2010). The mechanism of *pry-1*-mediated VPC induction and cell fate specification are not well understood.

To investigate the genetic network of *pry-1*, we focused on differentially expressed genes that are associated with reproductive structure development. Of the 149 candidates present in *pry-1* mutant transcriptome (Ranawade *et al*. 2018), we specifically investigated 26 genes that were overexpressed in mutant animals. RNAi knockdown experiments revealed that eight of these (*spp-1, clsp-1, ard-1, cdk-1, rpn-7, his-7, rnr-1*, and *cpz-1*) strongly suppressed the *pry-1* Muv phenotype. Except for *spp-1* and *clsp-1* all others also inhibited VPC induction defects in *pry-1* mutants. We examined genetic interactions of the genes with *bar-1* in vulval cells, which revealed that a subset *(ard-1, cdk-1, his-7, rnr-1, and cpz-1)* may act downstream of *pry-1-bar-1*-mediated signaling. All of the suppressor genes are conserved in eukaryotes and perform diverse functions such as oxidation-reduction reactions (dehydrogenase family: *ard-1* and reductase family: *rnr-1*), protein degradation (peptidase *cpz-1* and proteasomal complex component *rpn-7*), protein phosphorylation (kinase *cdk-1*), and regulation of gene expression (histone *his-7*). We investigated whether suppressors also participate in other *pry-1*-mediated processes, such as stress response and lifespan maintenance. Recent work from our lab has shown that *pry-1* utilizes *daf-16 (FOXO* family) to regulate the lifespan of animals (Mallick *et al*. 2020). RNAi knockdown of *cdk-1, his-7, rnr-1*, and *cpz-1* significantly rescued the stress sensitivity and short lifespan of *pry-1* mutants. Interestingly, *cdk-1* and *rnr-1* RNAi extended the lifespan of *daf-16* mutants, suggesting that these two genes negatively regulate *pry-1-* and *daf-16*-mediated aging. Additional genetic epistasis experiments revealed that *his-7* is likely to function upstream and *cpz-1* downstream or independent of *daf-16*. Overall, these findings demonstrate that PRY-1 interacts with a diverse set of conserved proteins to control essential biological processes in *C. elegans*.

## MATERIALS AND METHODS

### Strains

Animals were maintained at 20°C on standard nematode growth media (NGM) plates seeded with OP50 *E. coli* bacterial strains as described by Brenner (Brenner 1974). Worm strain information can be found in **Supplementary Table S1**.

### RNAi

RNAi mediated gene silencing was performed using a protocol previously published by our laboratory (Ranawade *et al*. 2018). Plates were seeded with *Escherichia coli* HT115 expressing either dsRNA specific to candidate genes or empty vector (L4440). Synchronized gravid adults were bleached, and eggs were plated. After becoming young adult animals were analyzed for vulva or seam cell phenotype.

### Vulva phenotype and VPC induction analysis

The mutivulva (Muv) and protruding vulva (Pvl) phenotypes were scored in adults at plate level. An animal with two or more protrusions (pseudovulvae) on its ventral side was assigned to a class of Muv mutant. Whereas an animal with a single prominent pseudovulva were placed in the Pvl class. VPC induction was determined at the mid-L4 stage. In a wild-type worm, the induction score is 3-one each for P5.p, P6.p and P7.p. Muv animals have the score of greater than 3 due to the induction of additional VPCs.

### Lifespan analysis

All lifespan analysis was done following adult specific RNAi treatment using a protocol described previously (Mallick *et al*. 2020). Animals were grown on NGM OP50 seeded plates till late L4 stage after which they were transferred to RNAi plates. Plates were then screened daily for dead animals and surviving worms were transferred every other day till the progeny production ceased. Censoring was done for animals that either escaped, burrowed into the medium, showed a bursting of intestine from the vulva or formed a bag of worms (larvae hatches inside the worm and the mother dies).

### Stress assay

Oxidative stress experiments were performed by exposing animals to 100mM paraquat (PQ) for 1h and 2h using a published protocol (Li et al 2008). Final working concentrations were made in M9 instead of water. At least 30 animals were tested for each strain in each replicate. Mean and standard deviation were determined from experiments performed in duplicate. Animals were considered dead if they had no response following a touch using the platinum wire pick and showed no thrashing or swimming movement in M9. Moreover, dead animals usually had an uncurled body posture compared to the normal sinusoidal shape of worms.

### Body bending and pharyngeal pumping analysis

Rate of body bending per one min and the rate of pharyngeal pumping per 30 sec for adults were analysed over the period of four days (Collins *et al*. 2008). Hermaphrodites were analysed for these phenotypes under the dissecting microscope in isolation rather than in groups on OP50 plates. Pharyngeal pumping was assessed by observing the number of pharyngeal contractions for 30 sec. For body bending assessment, animals were stimulated by tapping once on the tail of the worm using the platinum wire pick where one body bend corresponded to one complete sinusoidal wave of the worm. Only animals that moved throughout the duration of one min were included in the analysis.

### Molecular Biology

RNA was extracted from synchronized L3 and day-1 adult animals. Protocols for RNA extraction, cDNA synthesis and qPCR were described earlier (Ranawade *et al*. 2018). Briefly, total RNA was extracted using Trizol (Thermo Fisher, USA), cDNA was synthesized using the SensiFast cDNA synthesis kit (Bioline, USA), and qPCR was done using the SYBR green mix (Bio-Rad, Canada). Primers used for qPCR experiments are listed in **Supplementary Table S1**.

### Fluorescent microscopy

Animals were anaesthetized using 10mM Sodium Azide and mounted on glass slides with 2% agar pads and covered with glass coverslips. Images were captured using Zeiss Apotome microscope and Zeiss software. Fluorescence of *hsp-4::GFP, hsp-60::GFP, sod-3::GFP* and *daf-16p::DAF-16::GFP* was examined by analyzing the degree of GFP intensity. Quantification of pixel densities for GFP reporters was performed with ImageJ™ (https://imagej.nih.gov/ij/).

### Statistical analyses

Statistics analyses were performed using SigmaPlot software 11, CFX Maestro 3.1 and Microsoft Office Excel 2016. For lifespan data, survival curves were estimated using the Kaplan-Meier test and differences among groups were assessed using the log-rank test. qPCR data was analyzed using Bio-Rad CFX Maestro 3.1 software. For all other assays, data from repeat experiments were pooled and analyzed together and statistical analyses were done using GraphPad Prism 8. *p* values less than 0.05 were considered statistically significant.

## RESULTS

### Expression of reproductive structure development genes is affected in the *pry-1* mutant

We used GO enrichment analysis (http://geneontology.org/) to filter differentially expressed genes associated with ‘reproductive structure development’ (GO:0048608) in the *pry-1* mutants and identified 149 genes (gene_ontology.obo file pulled from version control on May 3 08:44:20 2017 UTC and WormBase release WS258) (**Supplementary Table S2**). Among them, 52 and 97 genes were upregulated and downregulated, respectively, in the *pry-1* mutant transcriptome (Ranawade *et al*. 2018) (**Figure 1, A and B, Supplementary Table S2**). GO term analysis was performed to identify the genes associated with biological functions, molecular functions, and cellular localization. Significant enrichment (FDR *p* < 0.05) was observed for processes such as cellular component organization or biogenesis (69), anatomical structure development (61), metabolic process (57), regulation of transcription (27), cell cycle (24), and nervous system development (21) (**Supplementary Table S3**). Among the molecular functions, we observed enrichment for categories such as protein binding activity (64), DNA binding activity (27), RNA binding activity (20), hydrolase activity (26), signaling receptor binding activity (8), and protein kinase binding activity (7) (**Supplementary Table S3**). Many genes were found to be associated with the cellular components, including the nucleus (74), protein-containing complexes (65), cytoplasm (52), integral components of the membrane (20), cytoskeleton (17), and nuclear chromosomes (12) (**Supplementary Table S3**). This suggests that *pry-1* plays an essential role in regulating the expression of diverse sets of genes.

**Figure 1:**
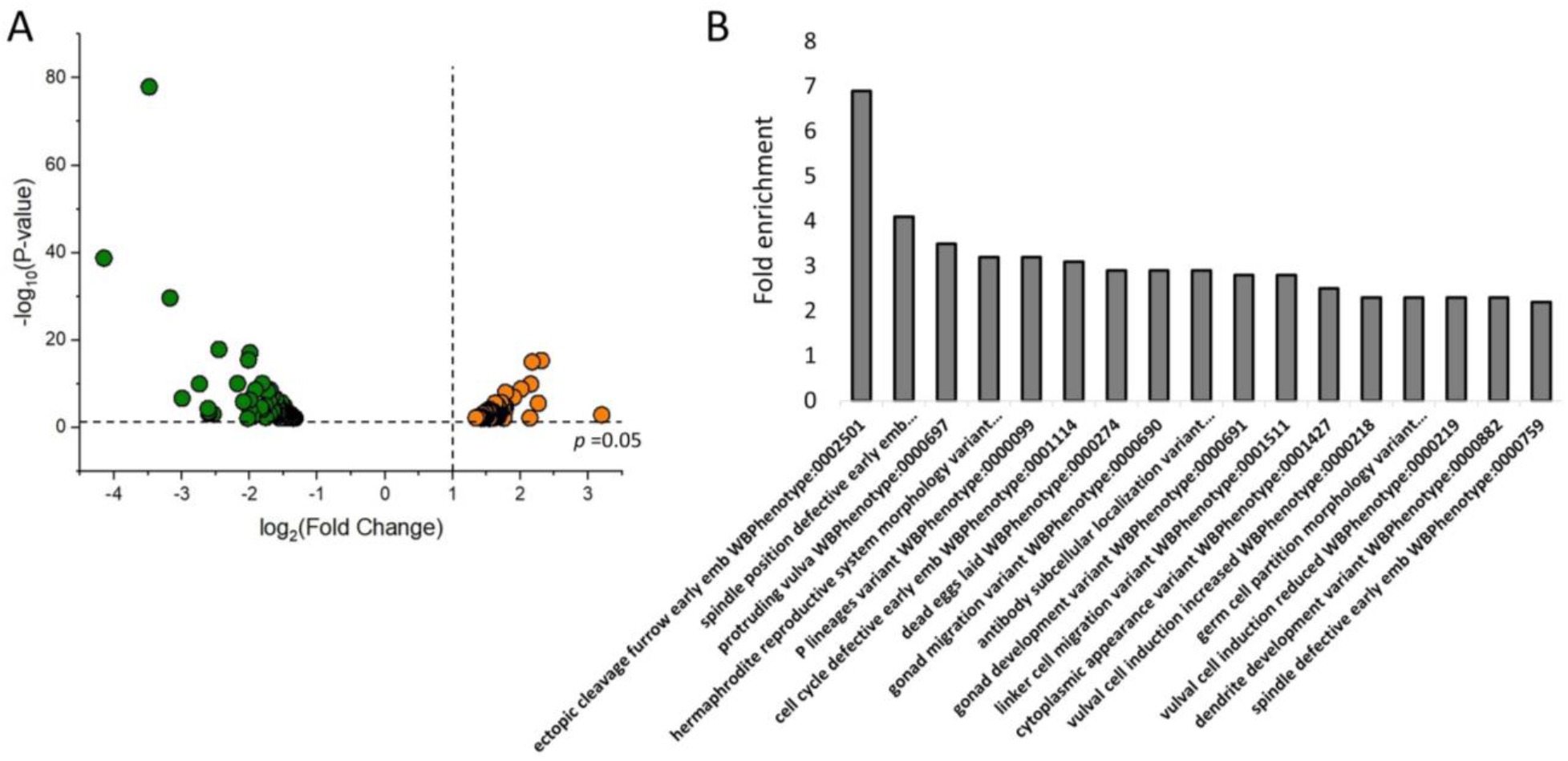
*pry-1* mutant transcriptome is enriched with genes involved in reproductive structure development. (A) Volcano plot showing the differentially expressed genes linked to reproductive structure development in *pry-1* mutant animals (**Supplementary Table S2**). (B) Phenotype-enrichment analysis of genes shown in (A). Not all categories are shown. See **Supplementary Table S4** for a complete list.

Tissue-enrichment and phenotype-enrichment analyses using a WormBase tool (https://wormbase.org/tools/enrichment/tea/tea.cgi, (Angeles-Albores *et al*. 2016)) indicated significant enrichment of tissues, such as neurons and P-cell lineages (**Supplementary Table S4**). The genes were also significantly associated with phenotypes such as protruding vulva, hermaphrodite reproductive system morphology variants, and vulval precursor cell induction increased or decreased (**Supplementary Table S4**).

### RNAi knockdown of a subset of reproductive structure development genes suppressed *pry-1* Muv phenotype

We evaluated the role of the upregulated genes in *pry-1*-mediated vulval development. Among the 52 upregulated genes, 26 (50%) were experimentally tested by RNAi to determine their effect on the Muv phenotype of *pry-1(mu38)* animals (**Figure 2**). Adult *pry-1* mutant hermaphrodites exhibit ectopic pseudovulvae-like structures. This is attributed the inappropriate induction of VPCs, resulting in the Muv phenotype in adults, in response to the constitutive activation of the canonical WNT signaling (Gleason *et al*. 2002). Sixteen genes significantly suppressed the Muv defect at the plate level (*p* < 0.05) (**Figure 2**). For eight of the genes, the reduction in vulva phenotype was lower than the mean +/-2□ of control RNAi-treated animals (**Figure 2**); therefore, we designated them as *pry-1* suppressors.

**Figure 2:**
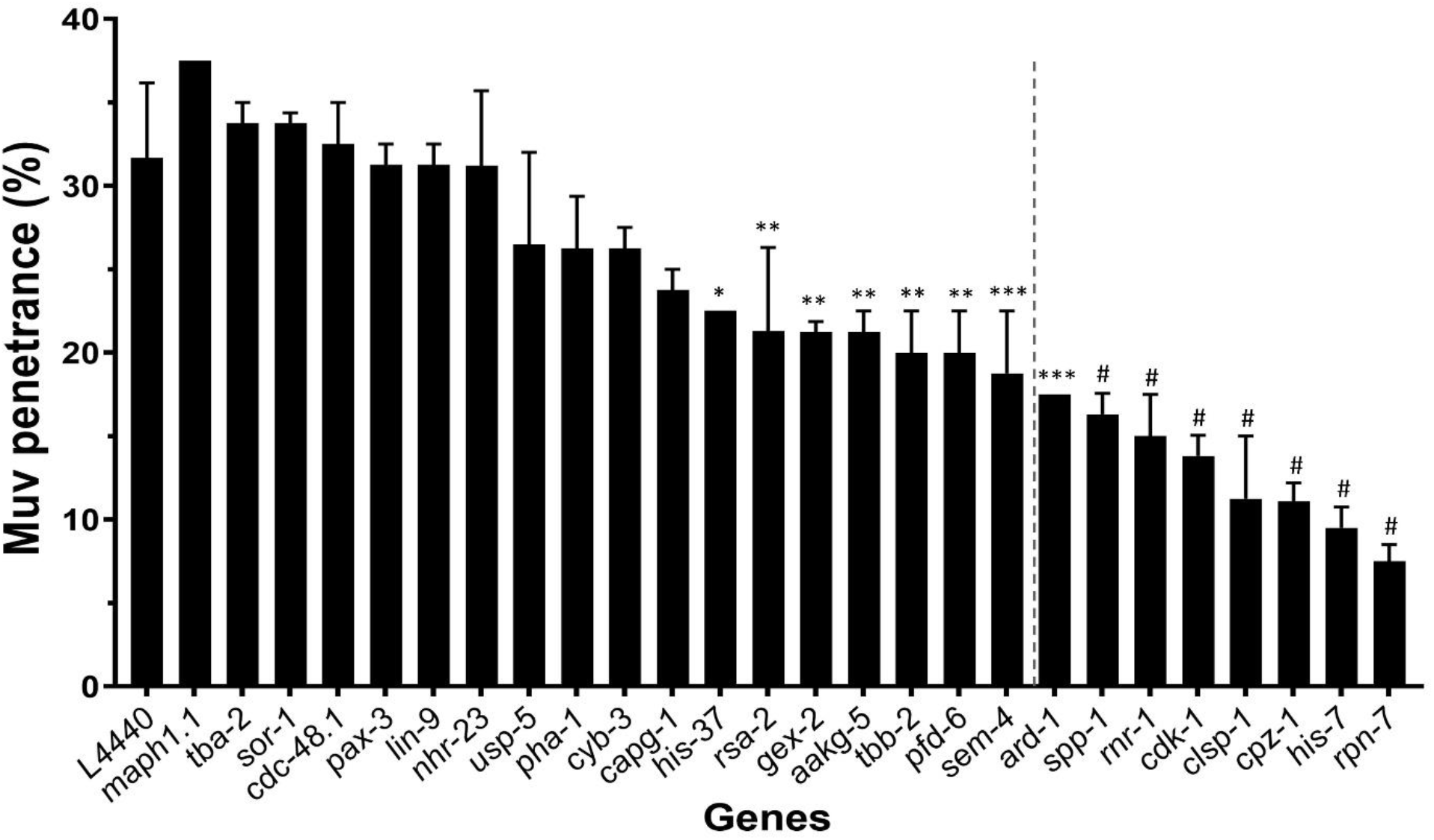
Quantification of multivulva phenotype following RNAi knockdown of 26 upregulated genes in *pry-1(mu38)* animals. Data represents the mean of two replicates (n > 40 animals in each replicate) and error bars represent the standard deviation. Genes on the right of the dotted vertical line were investigated further. Statistical analyses were done using one-way ANOVA with Dunnett’s post hoc test and significant differences are indicated by stars (*): * (*p* <0.05), ** (*p* <0.01), *** (*p* < 0.001), # (*p* < 0.0001).

All the eight genes, except one (*spp-1*), have homologs in higher eukaryotes including humans, suggesting their important roles in conserved biological processes. Four of the suppressors encode proteins that possess or regulate enzymatic activities, specifically acting as oxidoreductases (*ard-1* and *rnr-1*), protease (*cpz-1*), and regulatory subunit of a proteasome complex (*rpn-7*). Two of the suppressors, *cdk-1* and *clsp-1*, are involved in protein phosphorylation. *cdk-1* is possibly necessary for all cell divisions in *C. elegans* (Boxem 2006). *his-7* is a member of the human H2A family of histones that function in DNA repair and gene expression, and *spp-1* is a homolog of the human gene encoding Saposin-like protein that plays a role in immunity (**Table 1**).

**Table 1:** List of eight suppressor genes, their mammalian homologs, associated GO-biological processes, and functions in *C. elegans*.

| Gene | Mammalian homolog | GO-biological process | Function in *C. elegans* |
| --- | --- | --- | --- |
| *ard-1* | Human HSD17B10 (hydroxysteroid 17-beta dehydrogenase 10) | Oxidation-reduction process | Dehydrogenase with NADP binding domain |
| *cdk-1* | Human CDK1 (cyclin dependent kinase 1) | Cell cycle; cell division; and protein phosphorylation | Serine/threonine kinase activity |
| *clsp-1* | Human CLSPN (claspin) | Mitotic G2 DNA damage checkpoint; activation of protein kinase activity; and mitotic DNA replication checkpoint | Anaphase-promoting complex binding activity; part of the Ataxia telangiectasia and Rad3-related (ATR)/ATL-1 DNA replication checkpoint pathway |
| *cpz-1* | Human CTSZ (cathepsin Z) | Vulval development; proteolysis; embryo development ending in birth or egg hatching; gonad morphogenesis; ecdysis; collagen and cuticulin-based cuticle | Predicted to have cysteine type endopeptidase activity |
| *his-7* | Human H2AX (histone H2A type 2-B) | DNA repair; chromatin organization; chromatin silencing; and regulation of transcription | Predicted DNA binding activity |
| *rnr-1* | Human RRM1 (ribonucleotide reductase catalytic subunit M1) | DNA replication; metabolic process; deoxyribonucleotide biosynthetic process; and oxidation-reduction process | Ribonucleoside-diphosphate reductase activity |
| *rpn-7* | Human PSMD6 (proteasome 26S subunit, non-ATPase 6) | Proteasome-mediated ubiquitin-dependent protein catabolic process | Proteasome 26S complex component; ATP-dependent degradation of ubiquitinated proteins |
| *spp-1* | Human SAPLIPs (saposin-like proteins, saposin-B type domain containing protein) | Pathogenesis; innate immune response; defense response to Gram-negative/positive bacterium; | Channel activity |
|  |  | and transmembrane transport |  |

To investigate whether increased expression of the suppressor genes contributes to the Muv phenotype in *pry-1(mu38)*, we performed qPCR experiments. Expression levels at the L3 stage when VPCs undergo division to produce vulval progeny were examined. All genes, except *cpz-1*, were upregulated (**Figure 3A**). In young adults most genes continue to be expressed at significantly high levels, except for *rpn-7* and *ard-1* (no change), and *his-7* (downregulated) (**Figure 3B**). These results, along with the developmental pattern analysis (**Supplementary Figure 1**), support the key roles of suppressor genes in vulva formation. In addition, the genes are likely to participate in other post-developmental processes.

**Figure 3:**
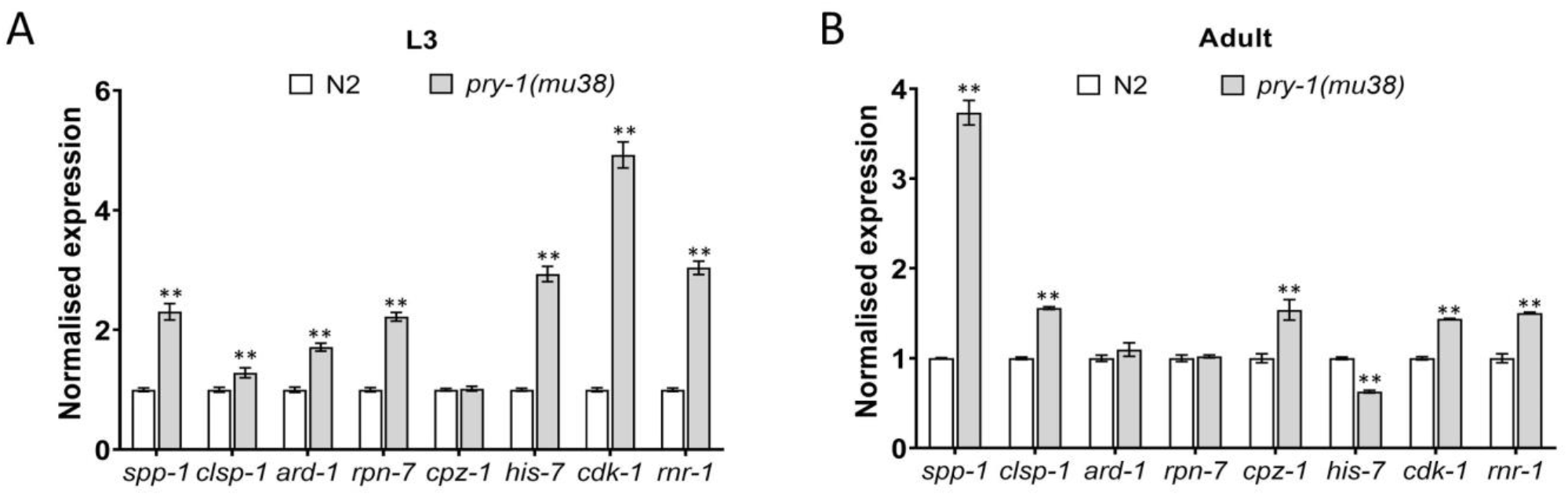
Expression levels of suppressor genes determined by qPCR in L3 and adults of *pry-1* mutants. Each data represents the mean of two replicates and error bars the standard error of means. Significance was calculated using Bio-Rad software (one-way ANOVA) and significant differences are indicated by stars (*): ** (*p* <0.01).

To understand the cellular basis of Muv suppression, the VPC induction pattern was investigated. RNAi of six of the eight suppressor genes strongly suppressed ectopic VPC induction in *pry-1* mutant animals. These include *ard-1, rpn-7, cpz-1, his-7, cdk-1*, and *rnr-1. cdk-1* and *rnr-1* RNAi induced the strongest suppression, with VPC induction being reduced by 46.7% and 50.2%, respectively (**Figure 4, A-C**) Interestingly, *clsp-1* and *spp-1* RNAi did not significantly suppress VPC induction, though both of them suppressed the *pry-1* Muv phenotype (**Figure 4, A-C**) One possibility may be that these two genes influence certain steps of morphogenesis thereby causing a reduction in pseudovulvae in *pry-1* mutants.

**Figure 4.**
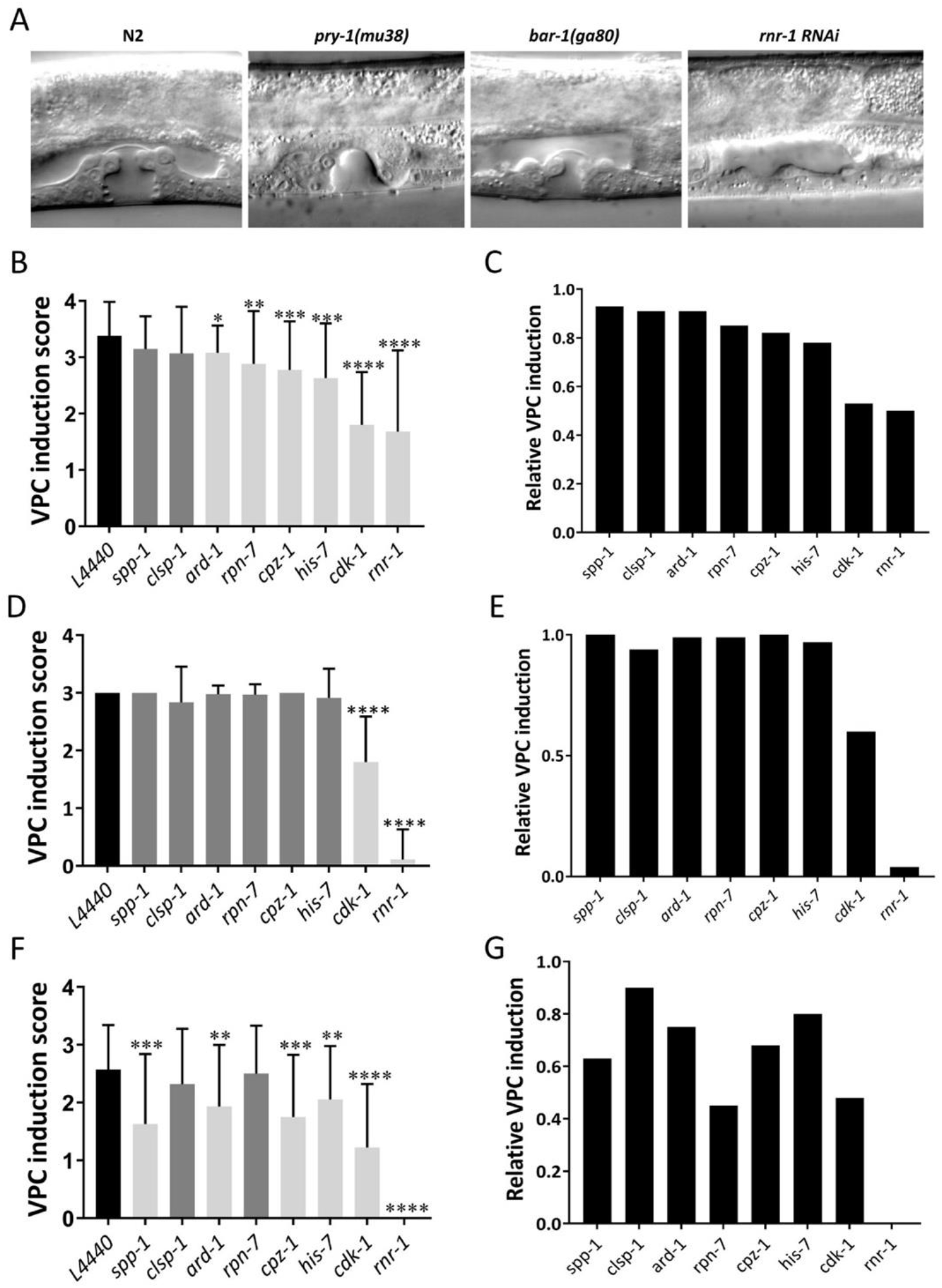
VPC induction analysis following RNAi knockdown of suppressor genes. (A) Representative image of N2, *pry-1(mu38), bar-1(ga80)* and *rnr-1* RNAi animals at the mid-L4 stage. (B) Knockdown of *ard-1, cdk-1, cpz-1, his-7, rnr-1* and *rpn-7* significantly reduced VPC induction in *pry-1(mu38)* animals. (C) Data in B plotted as a bar graph to depict changes in VPC induction. (D) Knockdown of *cdk-1* and *rnr-1* significantly reduced VPC induction in *N2* animals. (E) Data in D plotted as a bar graph to depict changes in VPC induction. (F) Knockdown of *ard-1, cdk-1, cpz-1, his-7, rnr-1* and *spp-1* significantly reduced VPC induction in *bar-1(ga80)* animals. (G) Data in F plotted as a bar graph to depict changes in VPC induction. In all cases, data represent a cumulative of two replicates (n > 30 animals in total for each condition) and error bars represent the standard deviation. Statistical analyses for panels B, D and F were done using one-way ANOVA with Dunnett’s post hoc test and significant differences are indicated by stars (*): ** (*p* <0.01), *** (*p* < 0.001), ****(*p*<0.0001)

We also examined the effects of suppressor genes on wild-type vulval development. RNAi experiments indicated no significant reduction in VPC induction for any of the genes except *cdk-1* and *rnr-1* (**Figure 4, A, D and E**). *cdk-1* and *rnr-1* RNAi resulted in 39% and 96.3% reduction in VPC induction, respectively, when compared to that in controls. These results along with those involving *pry-1* suggest that *cdk-1* and *rnr-1* possibly play essential roles in vulva formation and act genetically downstream of or in a pathway parallel to *pry-1*.

### Majority of the suppressor genes genetically interact with *bar-1/β-catenin*

PRY-1 is a component of the canonical WNT signaling pathway; therefore, we investigated whether any of the suppressor genes function as downstream effectors of the pathway during vulval development. To this end, we used the β-catenin homolog BAR-1, which is negatively regulated by PRY-1 (Eisenmann *et al*. 1998; Gleason *et al*. 2002). Mutations in *bar-1* cause some of the VPCs to remain uninduced. In a *bar-1* null mutant, *bar-1(ga80)*, P3.p and P4.p usually adopt an F (fused) fate. The frequency of F fate is much lower for the remaining VPCs (12–36% in each case), as they are mostly induced to form the vulva (Eisenmann *et al*. 1998; Eisenmann and Kim 2000). The VPC induction analysis following the RNAi knockdown of candidate genes revealed that *bar-1(ga80)* phenotype was significantly enhanced, for six of the genes, *ard-1, cpz-1, his-7, cdk-1, rnr-1*, and *spp-1. cdk-1* and *rnr-1* RNAi had the most severe effects, and the induction was reduced by 52% and 100%, respectively. In contrast, *clsp-1* and *rpn-7* did not significantly reduce VPC induction (**Figure 4, F and G**). These results, together with interaction experiments involving *pry-1*, suggest that *ard-1, cpz-1, his-7, cdk-1*, and *rnr-1* may act downstream of the *pry-1-bar-1* pathway to regulate VPC induction.

In addition to its role in VPC fate specification, *bar-1* is necessary for other aspects of vulva formation, because *bar-1(ga80)* adults exhibit a protruding vulva (Pvl) morphology. The Pvl phenotype is likely caused by abnormal fates of vulval progeny and their inability to invaginate correctly (Eisenmann and Kim 2000). We performed RNAi experiments to investigate whether suppressor genes influence the Pvl penetrance in *bar-1* mutants. The results revealed that *ard-1, his-7*, and *cdk-1* not only interacted with *bar-1* during VPC induction, but also participated in vulval morphogenesis (**Supplementary Figure S2**).

### Suppressor genes influence *pry-1*-mediated non-vulval processes

PRY-1 plays crucial roles in multiple developmental and post-developmental processes; therefore, we examined the involvement of Muv suppressors in all or subsets of PRY-1 non-vulval functions. The phenotypes examined include increased seam cell number, molting defect, low brood size, developmental delay, stress sensitivity, increased expression of chaperones (*hsp-4/BiP/GRP78, hsp-6/HSP70*, and *hsp-60/HSP60*), and short lifespan (Ranawade *et al*. 2018; Mallick *et al*. 2019a, 2020). Partial or complete loss of function of many of the suppressor genes is known to cause defects similar to *pry-1* mutants. These include *ard-1* RNAi leading to sterile progeny and developmental delay (Simmer *et al*. 2003; Sönnichsen *et al*. 2005), *cpz-1* and *rpn-7* RNAi animals showing molting defects (Hashmi *et al*. 2004; Frand *et al*. 2005), *clsp-1* RNAi leading to reduced germ line cell proliferation and increased expression of mitochondrial chaperones (Yoneda *et al*. 2004; Ceron *et al*. 2007), and *his-7* RNAi causing slow growth, larval arrest, and extended dauer survivability of *aak-2* mutants (Lehner *et al*. 2006; Ceron *et al*. 2007; Xie and Roy 2012).

To test whether the suppressor genes participate in *pry-1*-mediated developmental processes outside the vulva system, we first examined the seam cells. During seam cell development, *pry-1* promotes asymmetric cell division, and *pry-1* mutants show increased cell numbers. RNAi of the eight genes did not result in a change in the seam cells in both *pry-1* mutant and wild-type animals (**Supplementary Figure S3**), suggesting that none of these genes play a role in *pry-1*-mediated signaling in generating seam cells.

Next, we investigated whether suppressor genes interact with *pry-1* to regulate aging. Recent work from our group demonstrated the essential role of *pry-1* in mediating the lifespan maintenance in animals (Mallick *et al*. 2020). RNAi of the suppressor genes in *pry-1(mu38)* mutants from the L4 stage revealed that *cpz-1, his-7, cdk-1, and rnr-1* caused a significant extension in the mean lifespan (**Figure 5A and Table 2, Supplementary Figure S4**). We also observed that knockdown resulted in significant improvements in body bending and pharyngeal pumping, two physiological markers of aging (**Figure 5, C and D**). Interestingly, *cdk-1* and *rnr-1* RNAi had a beneficial effect on the wild-type genetic background (**Figure 5B and Table 2**), suggesting that these two genes play essential roles in restricting the lifespan of animals.

**Table 2:**
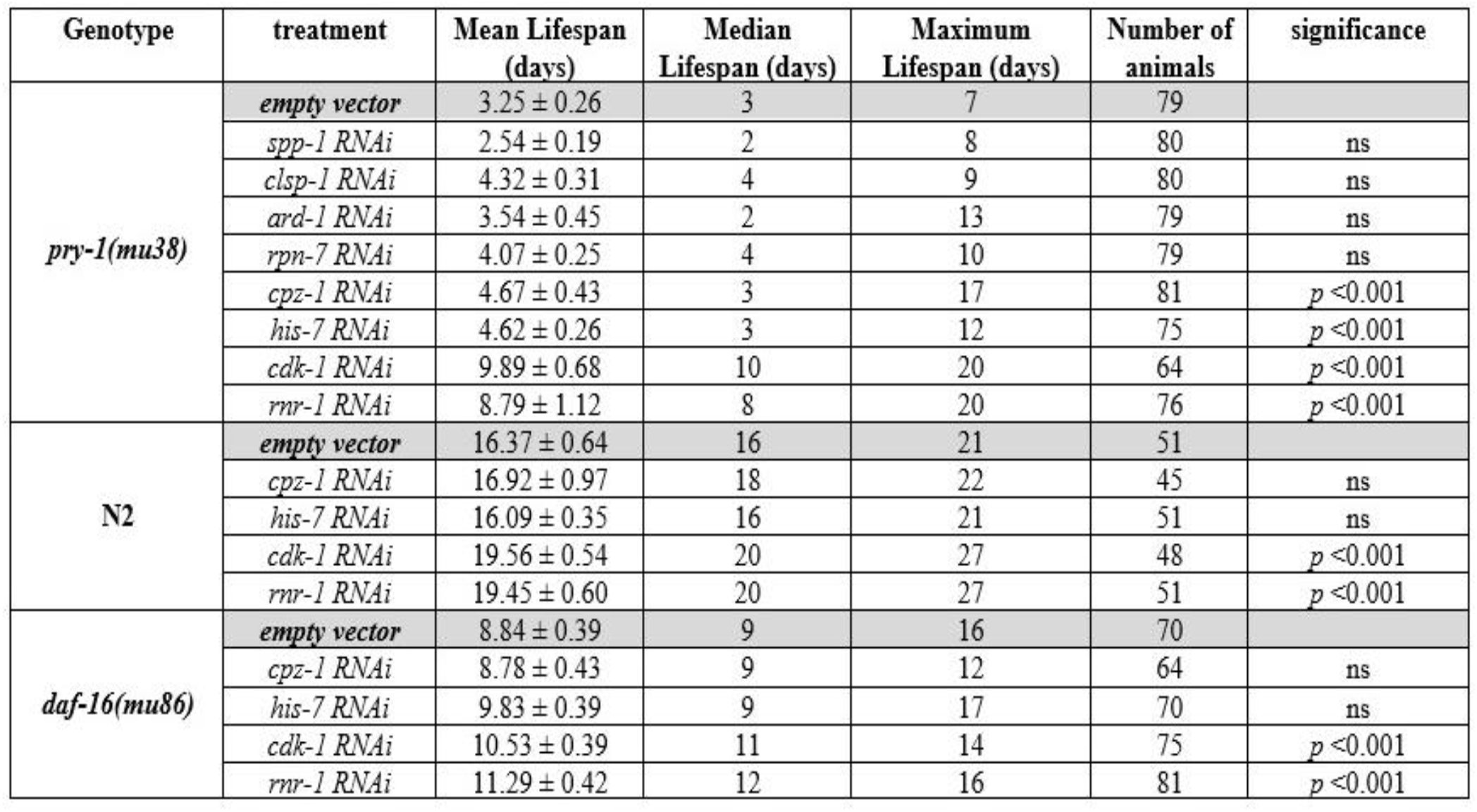
Mean, median, and maximum lifespan of N2, *pry-1(mu38)* and *daf-16(mu86)* animals following control (empty vector) and gene specific RNAi. In each case, data is presented as the cumulative of two replicates. ns: not significant.

**Figure 5:**
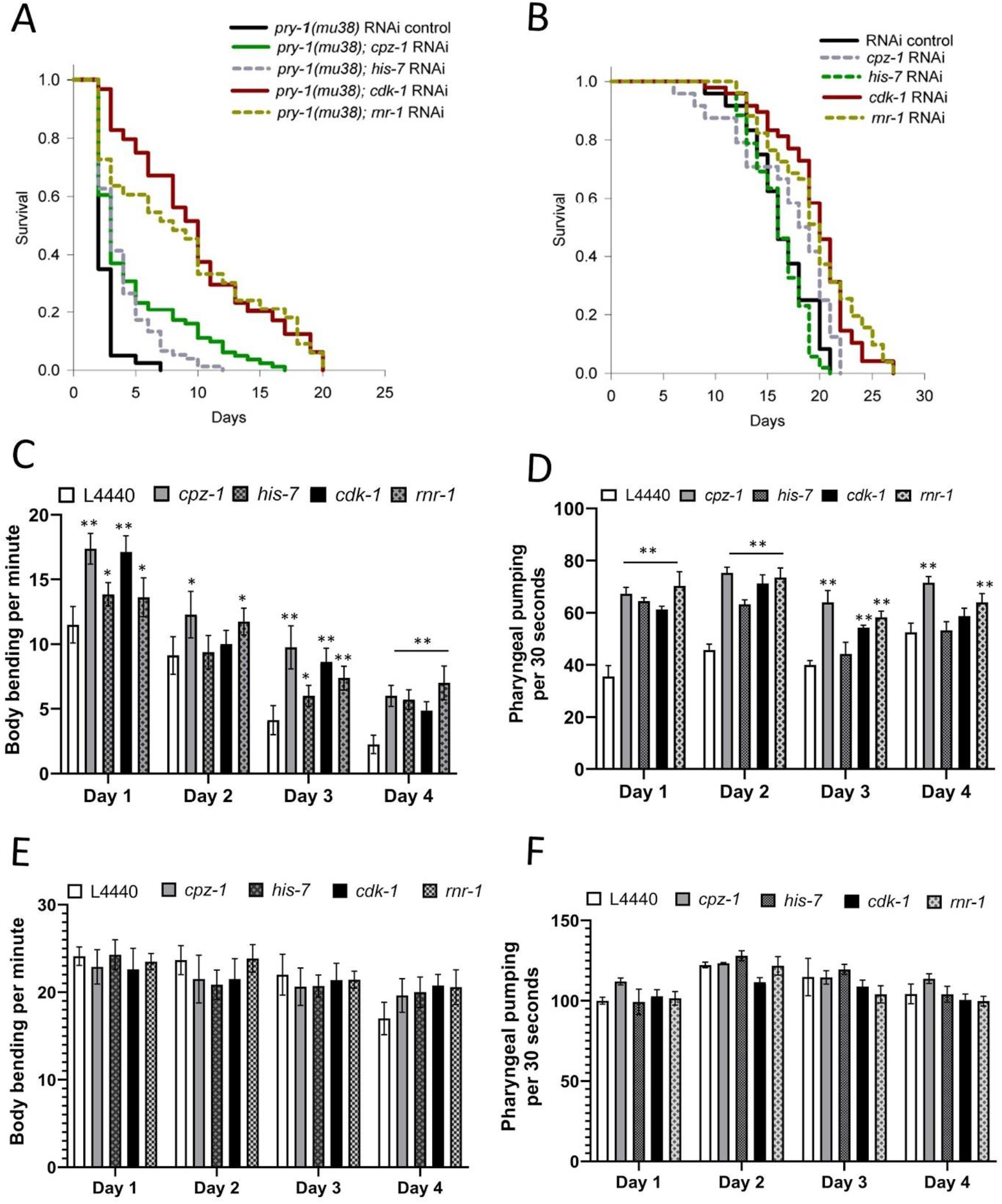
Knockdown of suppressor genes rescue lifespan, body bending and pharyngeal pumping defect of *pry-1* mutants. (A) RNAi of *cpz-1, his-7, cdk-1* and *rnr-1* extends lifespan of *pry-1* mutants (also see **Supplementary Figure S4**). (B) RNAi of *cdk-1* and *rnr-1* extends lifespan of N2 animals. (A-B) See Materials and Methods section and **Table 2** for lifespan data and statistical analyses. (C-D) Bar graphs showing rate of body bending and pharyngeal pumping of *pry-1* mutants over a period of 4 days following RNAi of *cpz-1, his-7, cdk-1 and rnr-1*. (E-F) Bar graphs showing rate of body bending and pharyngeal pumping of N2 animals. (C-F) Data represents the mean of two replicates (n >10 animals per replicate) and error bars represent the standard deviation. Statistical analyses for panels C-F were done using one-way ANOVA with Dunnett’s post hoc test for each day and significant differences are indicated by stars (*): * (*p* <0.05), ** (*p* <0.01).

In addition to its role in aging, *pry-1* is necessary for maintaining the expression of stress response signaling genes (Mallick *et al*. 2020). This led us to examine whether knockdown of suppressors lowers the lethality of *pry-1* mutants caused by acute exposure to the non-specific stress inducer PQ. Except for *clsp-1* and *rpn-7*, RNAi of the suppressor genes reduced the stress sensitivity in *pry-1* mutants (**Figure 6A**). Interestingly, *cpz-1* RNAi conferred stress resistance in wild-type animals (**Supplementary Figure S5**), suggesting the crucial role of this gene in stress response maintenance. Together with the lifespan analysis, these data suggest that *cdk-1, his-7, rnr-1*, and *cpz-1* contribute to *pry-1* mediated stress response and lifespan maintenance. The possibility that *cpz-1* acts in a pathway parallel to *pry-1* is also likely.

**Figure 6:**
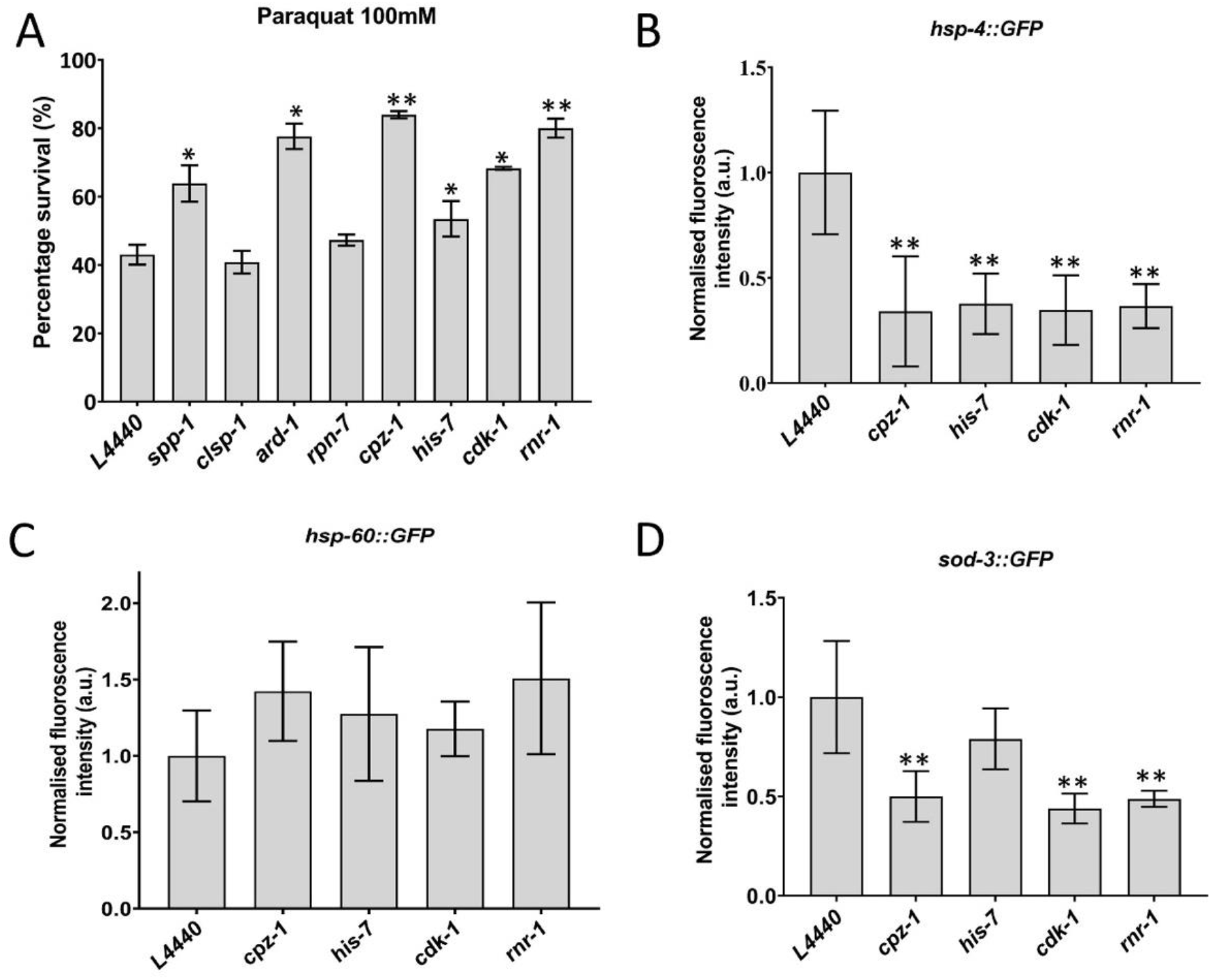
*cpz-1, his-7, cdk-1* and *rnr-1* regulate stress sensitivity in *pry-1* mutants. (A) Survivability of *pry-1(mu38)* animals following RNAi knockdown of genes. The animals were treated with 100mM PQ solution for one hour. Data represents mean of two replicates (n > 30 animals) and error bars represent standard deviation. (B) Quantification of fluorescence intensity using *hsp-4::GFP* marker in *pry-1* mutants following RNAi knockdown of *cpz-1, his-7, cdk-1* and *rnr-1*. (C) same as B, except that fluorescence reporter is *hsp-60::GFP* (D) same as B, except that fluorescence reporter is *sod-3::GFP*. Data represents the mean of two replicates (n >15 animals per replicate) and error bars represent the standard deviation. Statistical analyses for panels A-D were done using one-way ANOVA with Dunnett’s post hoc test for each day and significant differences are indicated by stars (*): * (*p* <0.05), ** (*p* <0.01).

We investigated whether downregulating the expression of *cdk-1, his-7, rnr-1*, and *cpz-1* in *pry-1* mutants could lower cellular stress (**Supplementary Figure S6, A and B**). *pry-1* mutants exhibit increased expression of three different stress response reporters: *hsp-4::GFP* (endoplasmic reticulum unfolded protein response chaperone), *hsp-60::GFP* (mitochondrial unfolded protein response chaperone), and *sod-3::GFP* (sodium dismutase, oxidative stress marker) (**Supplementary Figure S6B)**, in line with the previous report (Mallick *et al*. 2020). RNAi caused a significant suppression in *hsp-4::GFP* fluorescence (**Figure 6B**). A similar suppression in *sod-3::GFP* was observed, except for *his-7* (**Figure 6D**). There was no change in *hsp-60::GFP* fluorescence (**Figure 6C**). Overall, these results agree with our lifespan data and suggest that all the four suppressors are involved in multiple *pry-1*-mediated processes.

### Effect of *pry-1* suppressors on *daf-16/FOXO* lifespan phenotype

PRY-1-mediated lifespan regulation requires DAF-16 (Mallick *et al*. 2020); therefore, we investigated whether *cdk-1, his-7, rnr-1*, and *cpz-1* participate in the *pry-1-daf-16* genetic network. RNAi in *daf-16* mutant animals indicated that knockdown of *cdk-1* and *rnr-1* extended the lifespan (**Figure 7 and Table 2**). In contrast, RNAi of *his-7* and *cpz-1* had no such effect, consistent with the hypothesis that both of them act upstream of *daf-16*. There are two other possibilities: (1) the genes are downstream components but are insufficient on their own to rescue the *daf-16* mutant phenotype, and (2) they may act downstream of *pry-1* but in a *daf-16*-independent manner.

**Figure 7:**
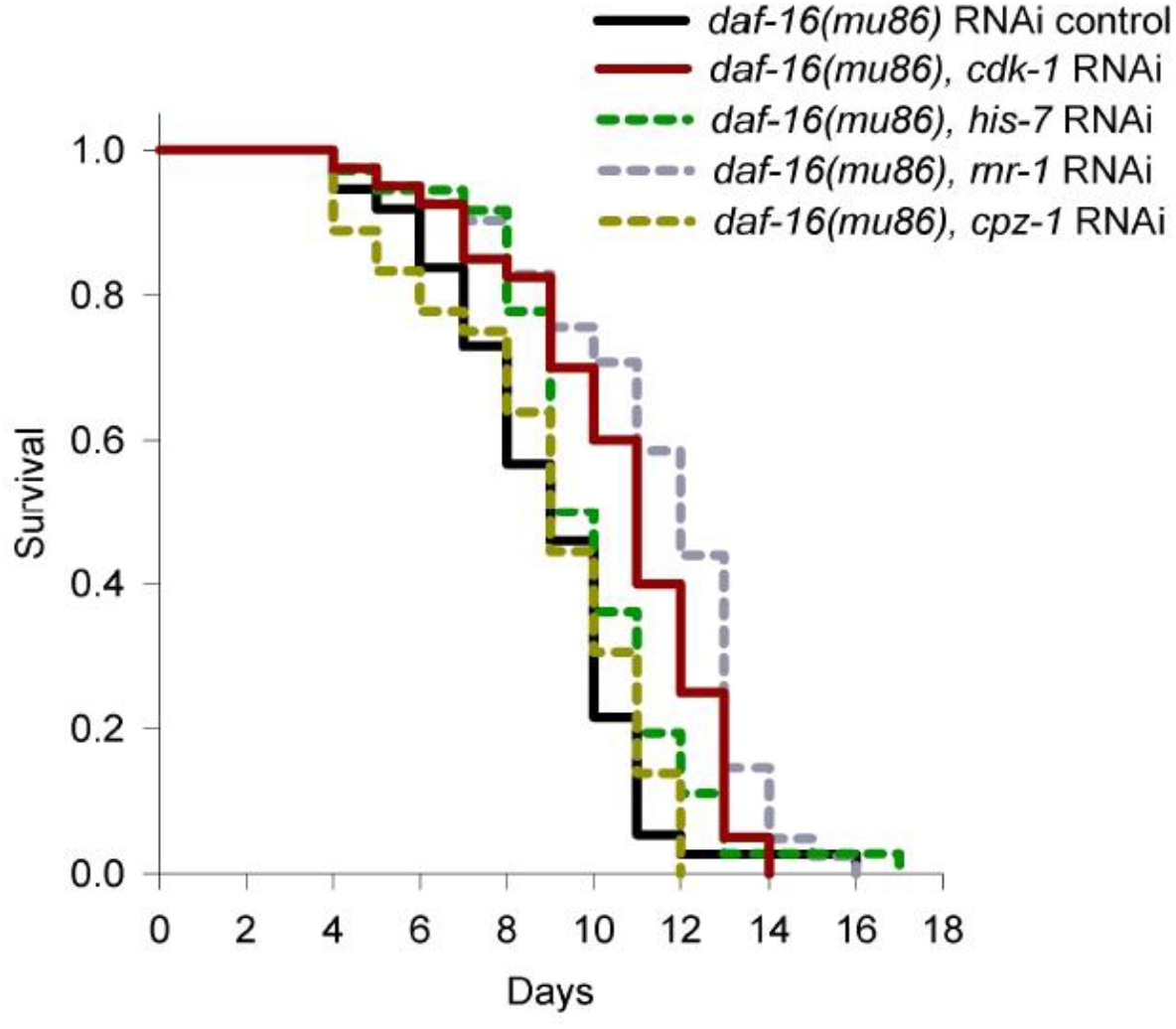
Lifespan analysis of *daf-16* mutants following RNAi knockdown of *cdk-1, his-7, rnr-1* and *cpz-1*. See Materials and Methods section and **Table 2** for lifespan data and statistical analyses.

To further investigate the genetic relationship of *his-7* and *cpz-1* with *daf-16*, we examined the fluorescence of DAF-16::GFP fusion protein, following RNAi. The localization of DAF-16::GFP was unaffected by both the genes; however, the fluorescence intensity was significantly reduced in *his-7* RNAi (**Supplementary Figure S7 and Table S5**). These results suggested that *his-7* might act genetically upstream, while *cpz-1* might function either downstream or independent of *daf-16*. It should be point out that *cpz-1* was reported to be upregulated in *daf-16* mutant animals (Lin *et al*. 2018). Further experiments are needed to determine the precise relationship between these two suppressor genes and *daf-16*. In conclusion, *rnr-1* and *cdk-1* negatively regulate the lifespan of wild-type animals, and *cpz-1* and *his-7* are not essential for normal lifespan; therefore, *cdk-1* and *rnr-1* may act downstream of the *pry-1-daf-16* pathway.

## DISCUSSION

In this study, we analyzed a set of genes that are upregulated in *pry-1* mutants and are associated with ‘reproductive structure development.’ The genes belong to GO categories, such as metabolic processes, transcriptional regulation, and mitotic cell cycle. Among the 26 genes tested, RNAi of eight suppressed the Muv phenotype of *pry-1* mutant animals with a threshold of mean +/-2□. Further analysis of VPC induction and genetic interactions with a null allele of *bar-1* revealed that five of the suppressors (*ard-1, cpz-1, his-7, cdk-1*, and *rnr-1*) are involved in *pry-1-bar-1*-mediated vulval cell proliferation. In addition, we found that *cdk-1* and *rnr-1* are essential for vulva development, because their downregulation caused defects in VPC induction in wild-type animals.

While our work provides the first evidence for the genetic interactions of *pry-1* with these eight genes (see schematic in **Figure 8**), some of the genes were previously reported to play roles in vulva formation. Specifically, disruption of *ard-1* and *clsp-1* causes a Pvl phenotype (Simmer *et al*. 2003; Ceron *et al*. 2007). *cpz-1* localizes to the developing vulva, and *cpz-1* RNAi results in defective vulval morphology (Hashmi *et al*. 2004). *cdk-1* regulates *lin-12/Notch* in a cell cycle-dependent manner (Nusser-Stein *et al*. 2012; Weinstein *et al*. 2015). The remaining four genes, *spp-1, rnr-1, rpn-7*, and *his-7* had no reported function in the vulva system.

**Figure 8:**
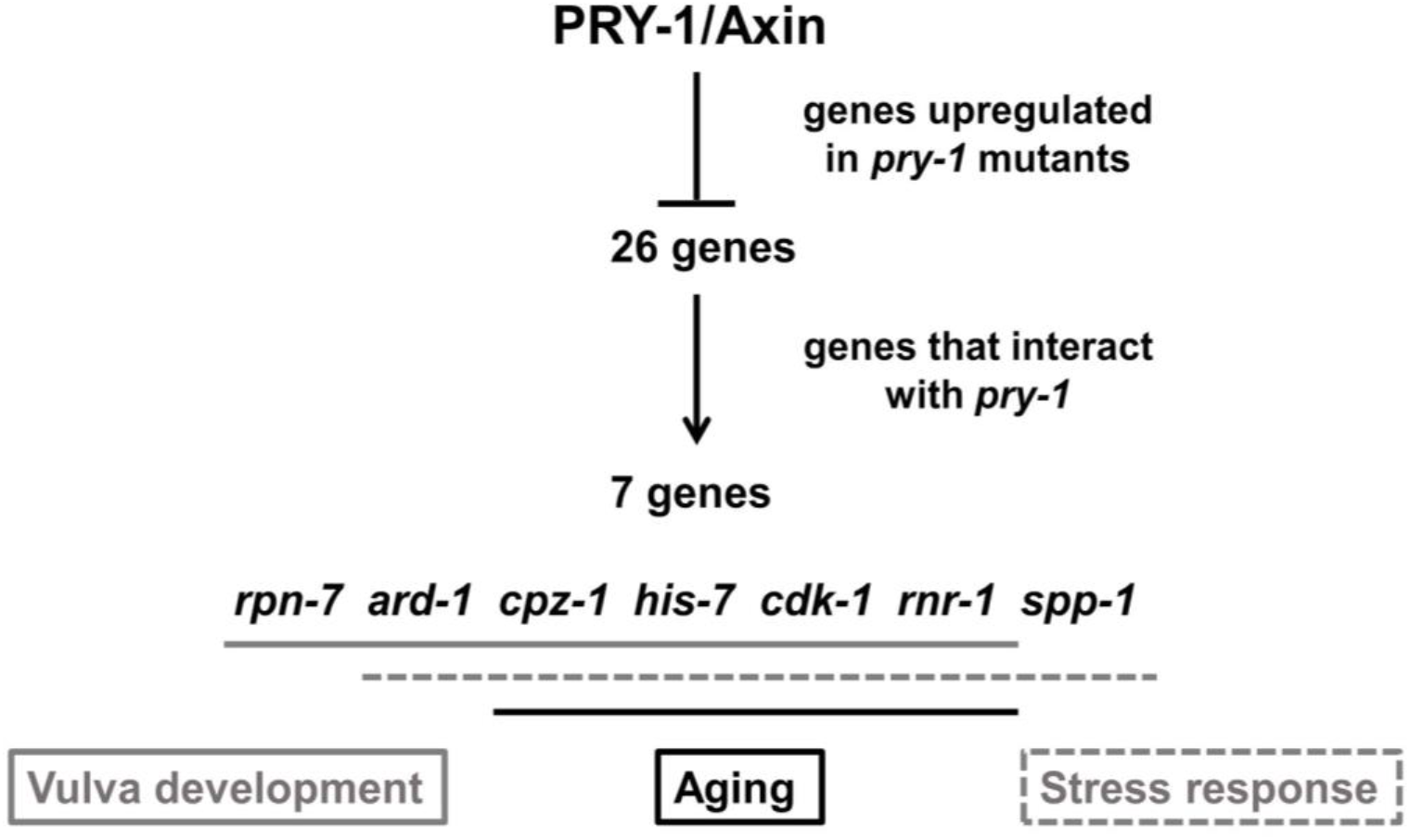
Schematic of genetic interactions of suppressor genes with *pry-1* in different processes. While *cpz-1, his-7, cdk-1* and *rnr-1* are involved in *pry-1*-mediated vulval development (grey solid line), aging (black solid line) and stress response (grey dotted line), *ard-1* plays a role in vulval development and stress response. *rpn-7* and *spp-1* interact with *pry-1* during vulval development and stress response, respectively.

In addition to studying vulval development, we investigated the role of the suppressor genes in *pry-1*-mediated post-developmental events. *cdk-1, rnr-1, rpn-7*, and *spp-1* are involved in aging. Specifically, *cdk-1* and *rpn-7* are both required for *glp-1*-mediated lifespan extension (Ghazi *et al*. 2007; Seidel and Kimble 2015), and *rpn-7* and *spp-1* are required for the longevity of *daf-2* mutants (Murphy *et al*. 2003; Ghazi *et al*. 2007; Anyanful *et al*. 2009). *rnr-1* is downregulated in long-lived *daf-2* and *eat-2* mutants, consistent with the notion that high levels of *rnr-1* decrease lifespan (Gao *et al*. 2018). These genes have potential roles in the conserved aging pathway; therefore, we analyzed their requirement in *pry-1* mediated lifespan regulation. *cdk-1, rnr-1, cpz-1*, and *his-7* function downstream of *pry-1*; they influence the *pry-1*-mediated stress response maintenance, as observed in the acute PQ assay and heat shock chaperone analysis. In addition, *his-7, cdk-1*, and *rnr-1* may be involved in the *daf-16*-mediated aging process. *his-7* is likely to act upstream of *daf-16*, while the other two genes may function downstream of the *pry-1-daf-16* pathway, for extending the lifespan of animals. All the four suppressor genes regulate fundamental cellular processes, such as protein phosphorylation (*cdk-1*), protein breakdown (*cpz-1*), DNA replication (*rnr-1*), and transcription (*his-7*) (**Table 1**). Consistent with this, *cdk-1* and *his-7* transcripts are enriched in germ line cells (Han *et al*. 2017), and *cpz-1* and *rnr-1* are expressed in the neurons, hypodermis, vulva, gonad, intestine, and muscles (Hashmi *et al*. 2004; Hunt-Newbury *et al*. 2007). Therefore, it is not surprising that perturbations in their function result in multiple phenotypes.

In conclusion, this study identified a new set of interactors of *pry-1* in *C. elegans*. Some of the suppressors affect multiple processes, while the others appear to have more restricted roles. All of the genes have mammalian orthologs, raising the possibility that their interactions with Axin may be conserved. Future studies hold promise to elucidate the mechanism by which these genes mediate tissue-specific function of Axin in normal and disease conditions.

## ACKNOWLEDGEMENTS

We thank Sakshi Mehta for assistance with certain experiments and Gupta lab members for discussions and suggestions. Some of the strains were obtained from CGC, which is funded by the NIH Office of Research Infrastructure Programs (P40OD010440).

## FUNDING

This work was supported by funds from the NSERC discovery grant to BPG.

## SUPPLEMENTAL MATERIALS

### FIGURES

**Figure S1:**
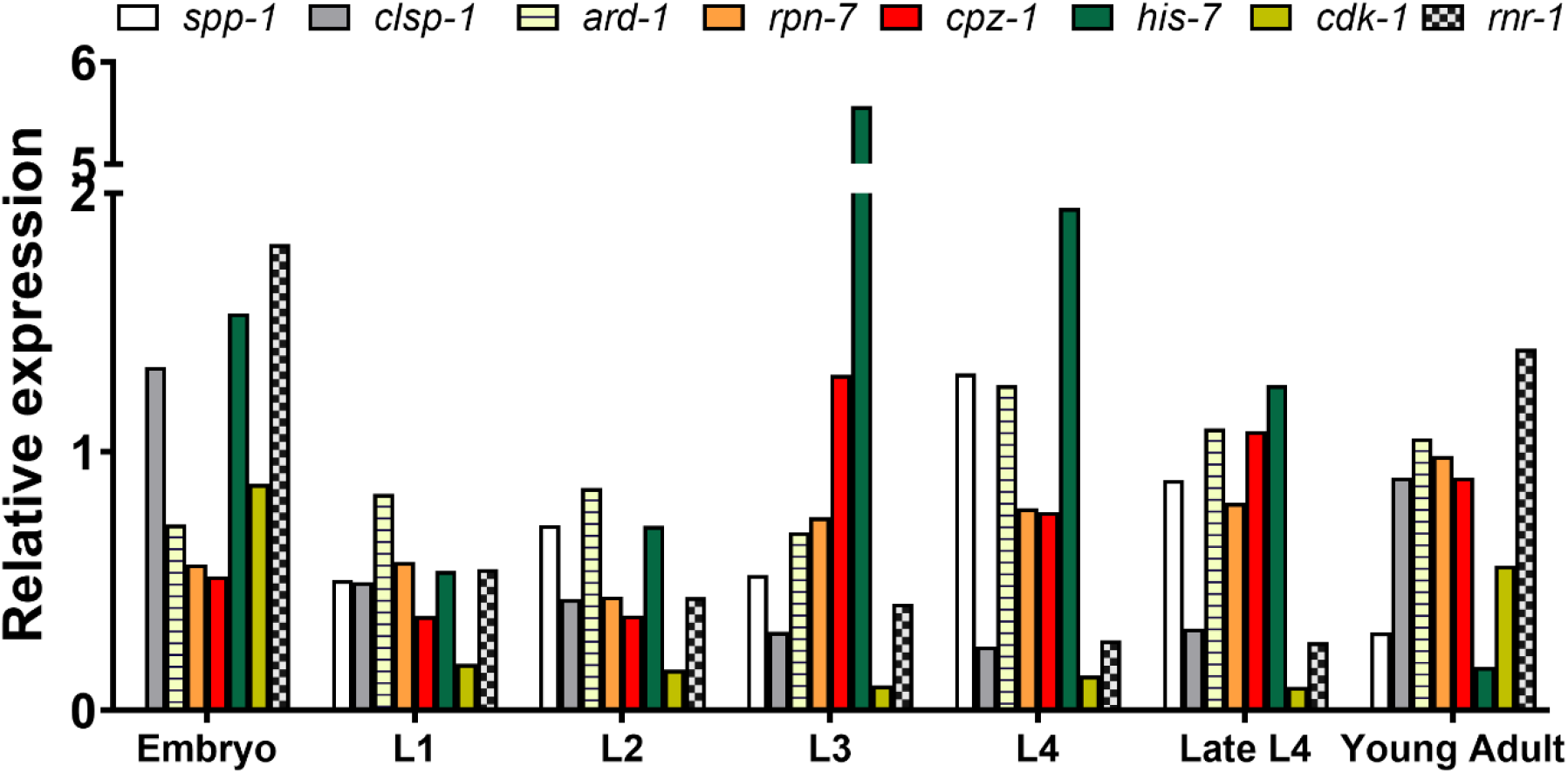
Expression of eight suppressor genes at different stages compared to mixed stage (reference), based on the previously reported RNAseq data (Grun et al 2014).

**Figure S2:**
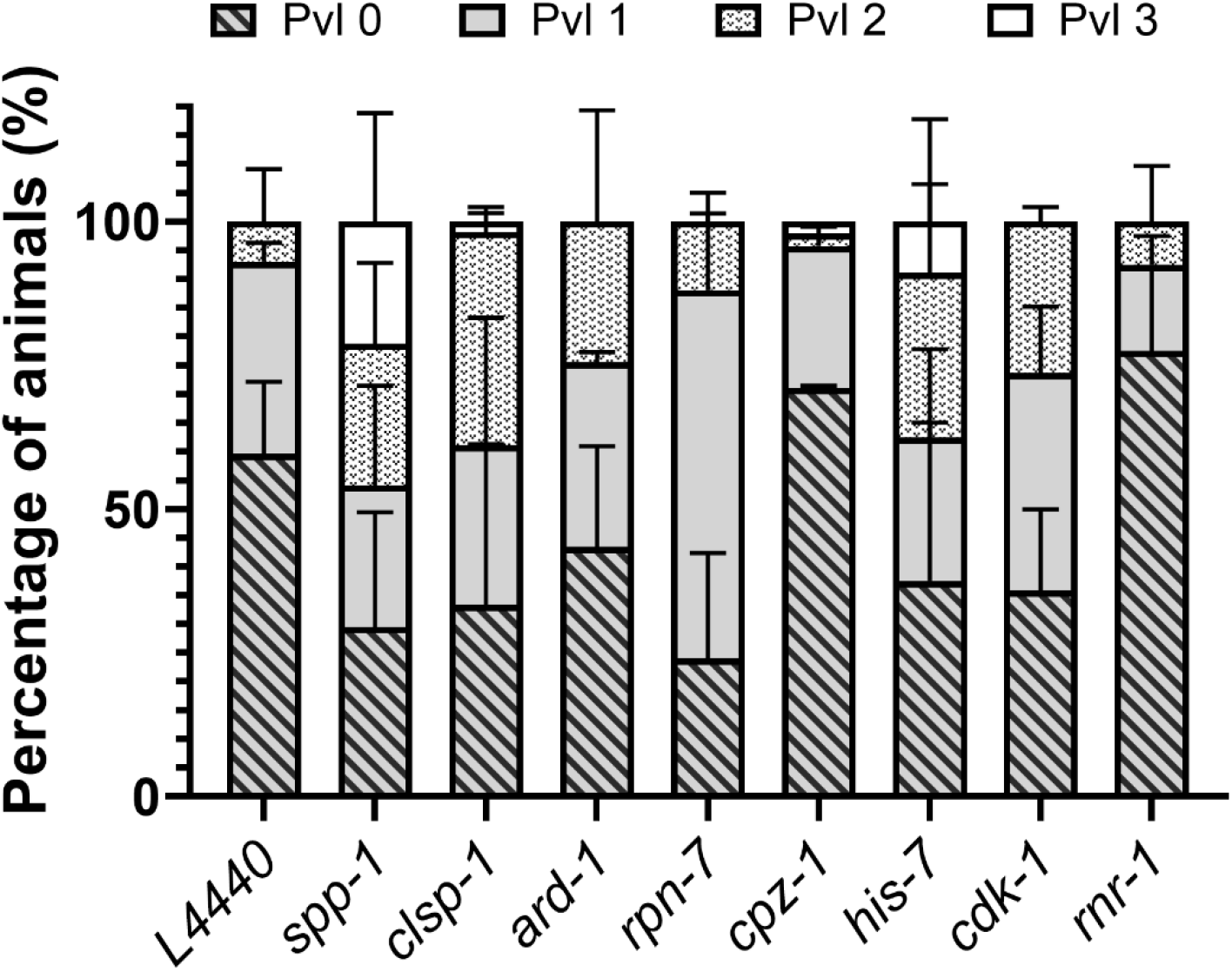
Protruding vulva analysis in *bar-1* mutants following RNAi knockdown of suppressor genes. The phenotype has been divided into 4 categories with Pvl 3 being the most severe. Data represents mean of two replicates (n > 30 animals) and error bars represent the standard deviation. Statistical analyses were done using two-way ANOVA with Tukey’s multiple comparison test and significant differences are indicated by stars (*): ****(*p*<0.0001). For Pvl 0, *spp-1, clsp-1, ard-1, his-7* and *cdk-1* RNAi is significantly different (****) to that of L4440; for Pvl 1, *rpn-7*, and *rnr-1* RNAi is significantly different (****) to that of L4440; for Pvl 2, *spp-1, clsp-1, ard-1, his-7*, and *cdk-1* RNAi is significantly different (****) to that of L4440; and for Pvl 3, only *spp-1* RNAi is significantly different (****) to that of L4440.

**Figure S3:**
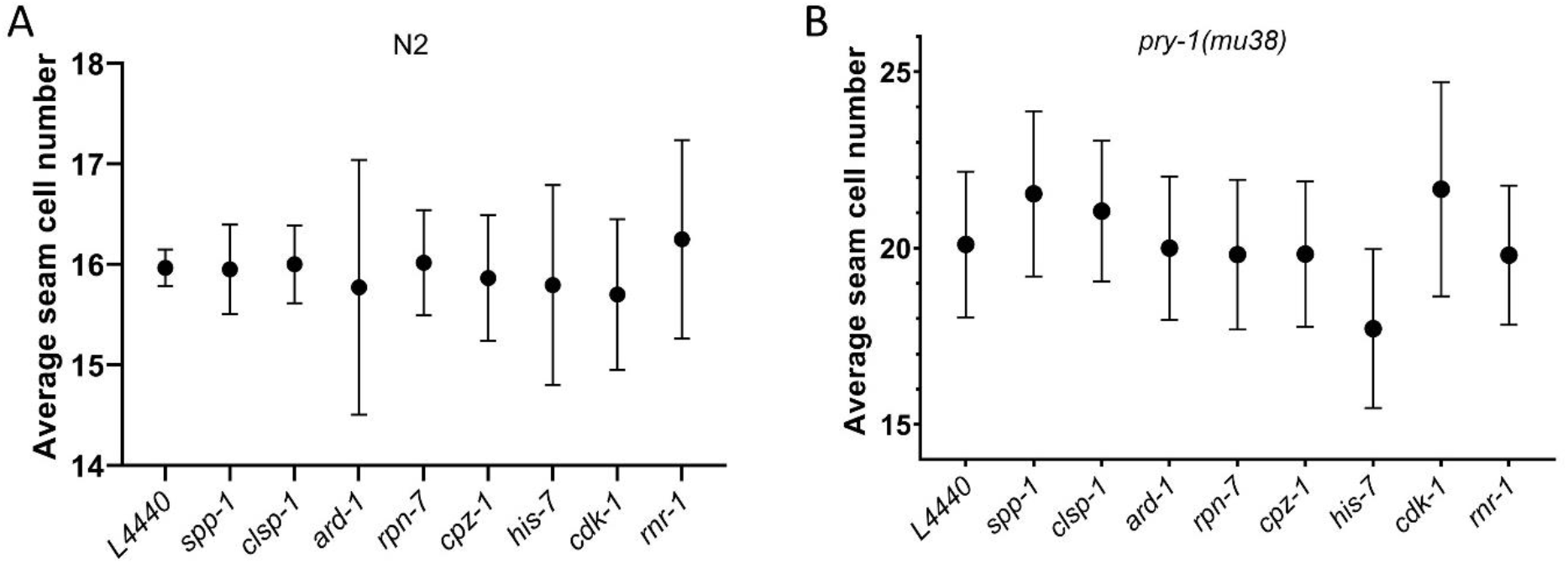
Seam analysis in both N2 and *pry-1(mu38)* animals following RNAi of control (L4440) and eight suppressor genes. Data represents mean of two replicates (n > 20 animals) and error bars represent the standard deviation. Statistical analyses were done using one-way ANOVA with Dunnett’s post hoc test.

**Figure S4:**
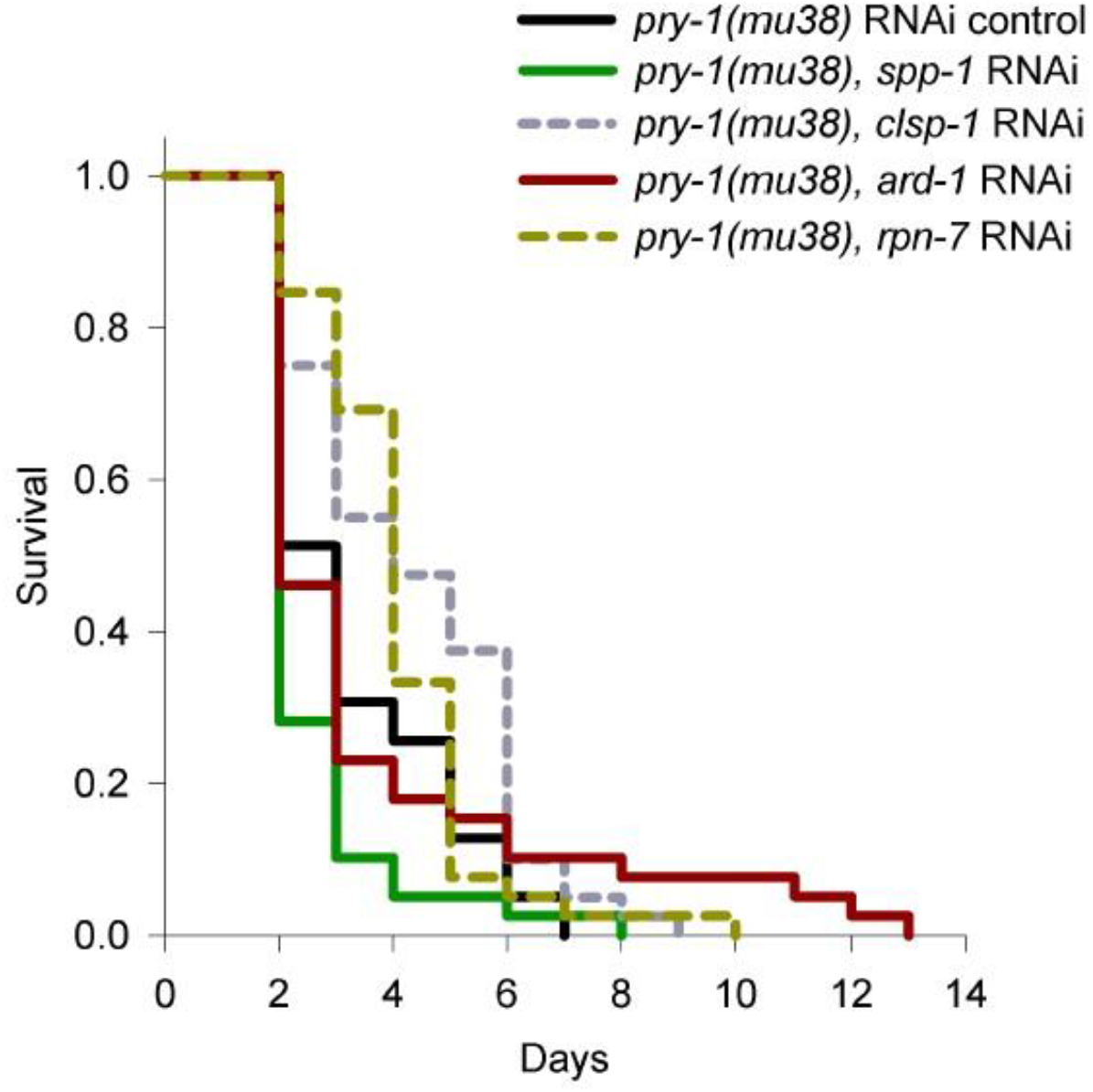
Lifespan analysis of *pry-1* mutants following RNAi knockdown of *spp-1, clsp-1, ard-1* and *rpn-7*. See Materials and Methods section and Table 2 for lifespan data and statistical analyses.

**Figure S5:**
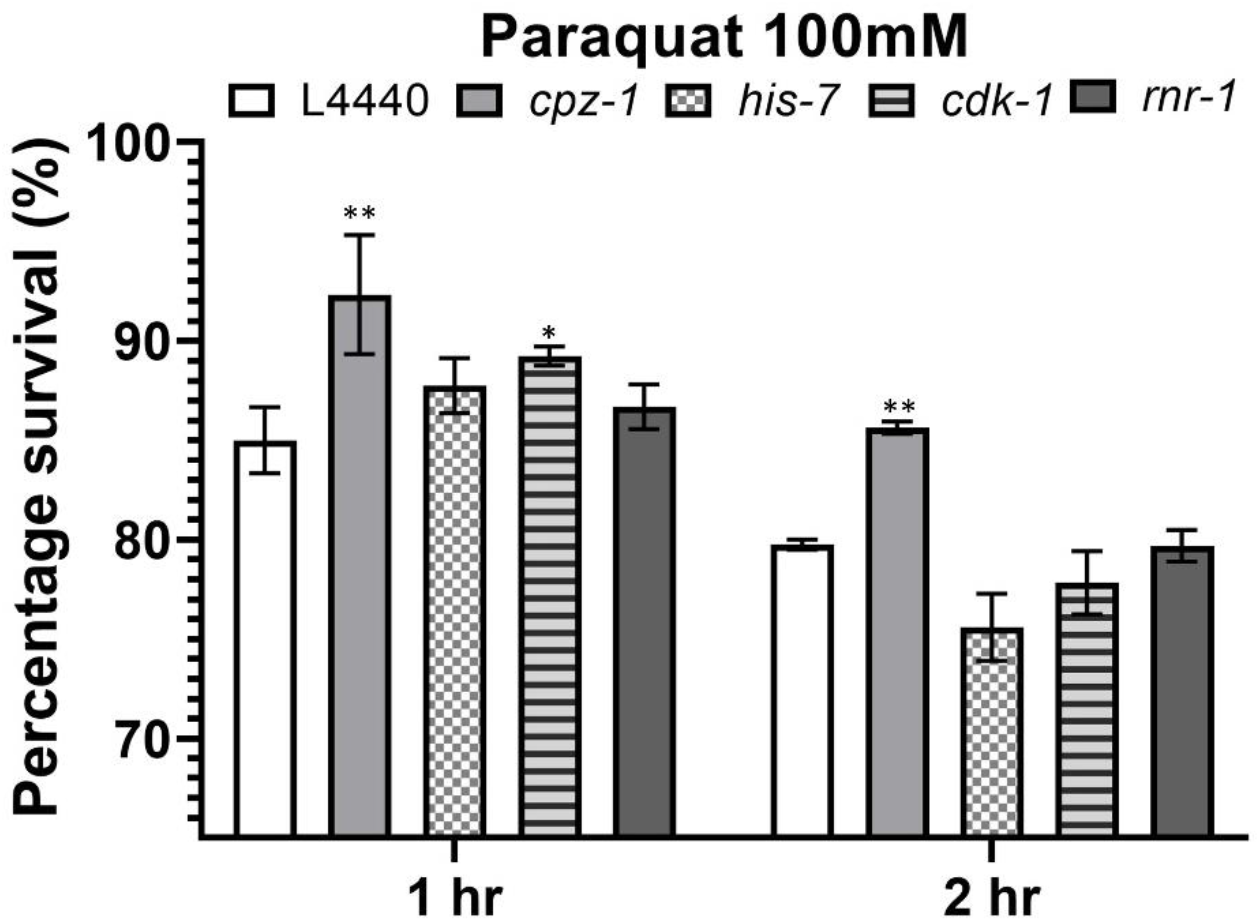
Survivability of N2 animals in 100mM PQ solution after 1 hour and 2 hour following RNAi knockdown of genes. Data represents mean of two replicates (n > 40 animals) and error bars represent the standard deviation. Statistical analyses were done using two-way ANOVA with Dunnett’s multiple comparison test for each hour and significant differences are indicated by stars (*): * (*p* < 0.05), ** (*p* <0.01).

**Figure S6:**
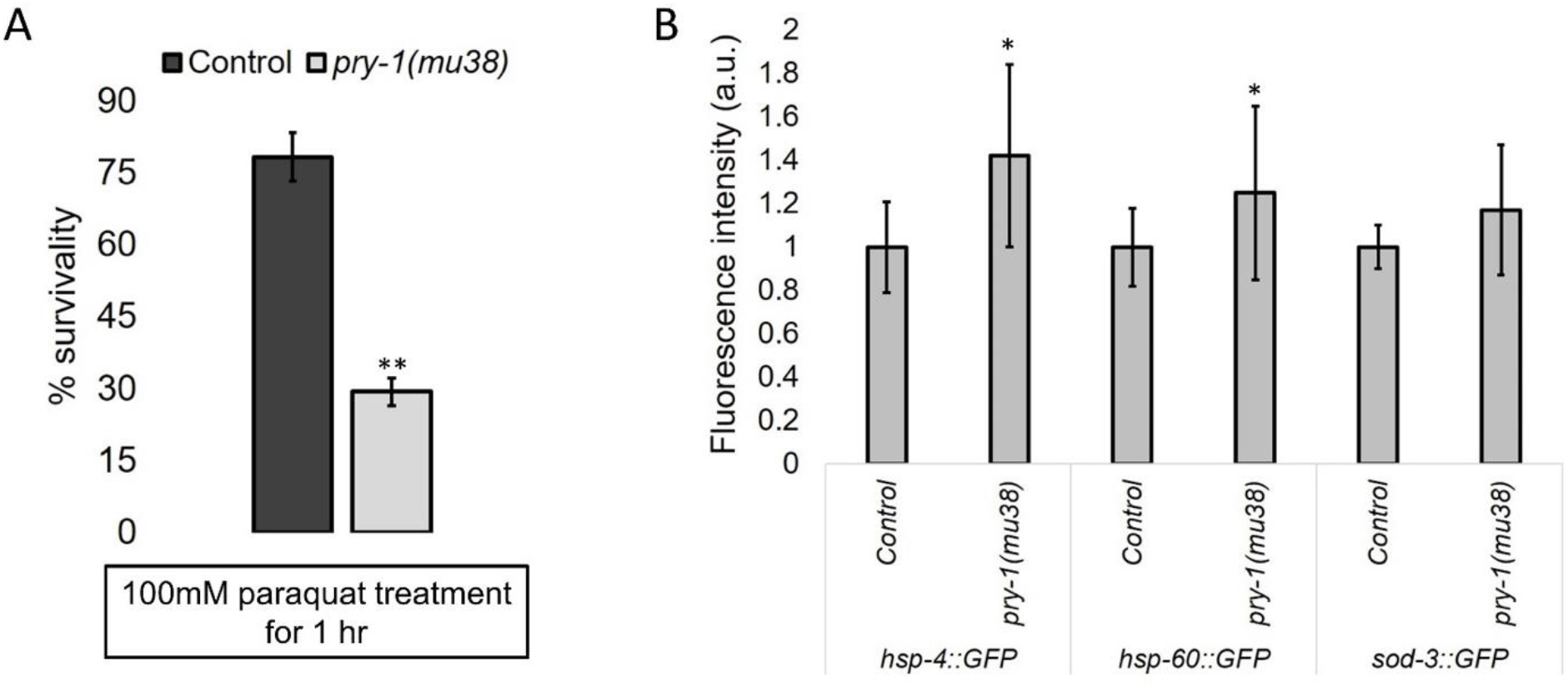
*pry-1* mutants are stress sensitive and show increased expression of stress response markers. (A) Stress sensitivity assay of control and *pry-1(mu38)* animals in 100mM PQ solution after 1hr. Data represents the mean of two replicates (n >30 animals per replicate) and error bars represent the standard deviation. (B) Quantification of fluorescence intensity using *hsp-4::GFP, hsp-60::GFP* and *sod-3::GFP* marker in control and *pry-1* mutants. Data represents the mean of two replicates (n >10 animals per replicate) and error bars represent the standard deviation. Statistical analyses for panels A and B were done using unpaired student’s t-test with unequal variance significant differences are indicated by stars (*): * (*p* < 0.05), ** (*p* <0.01).

**Figure S7:**
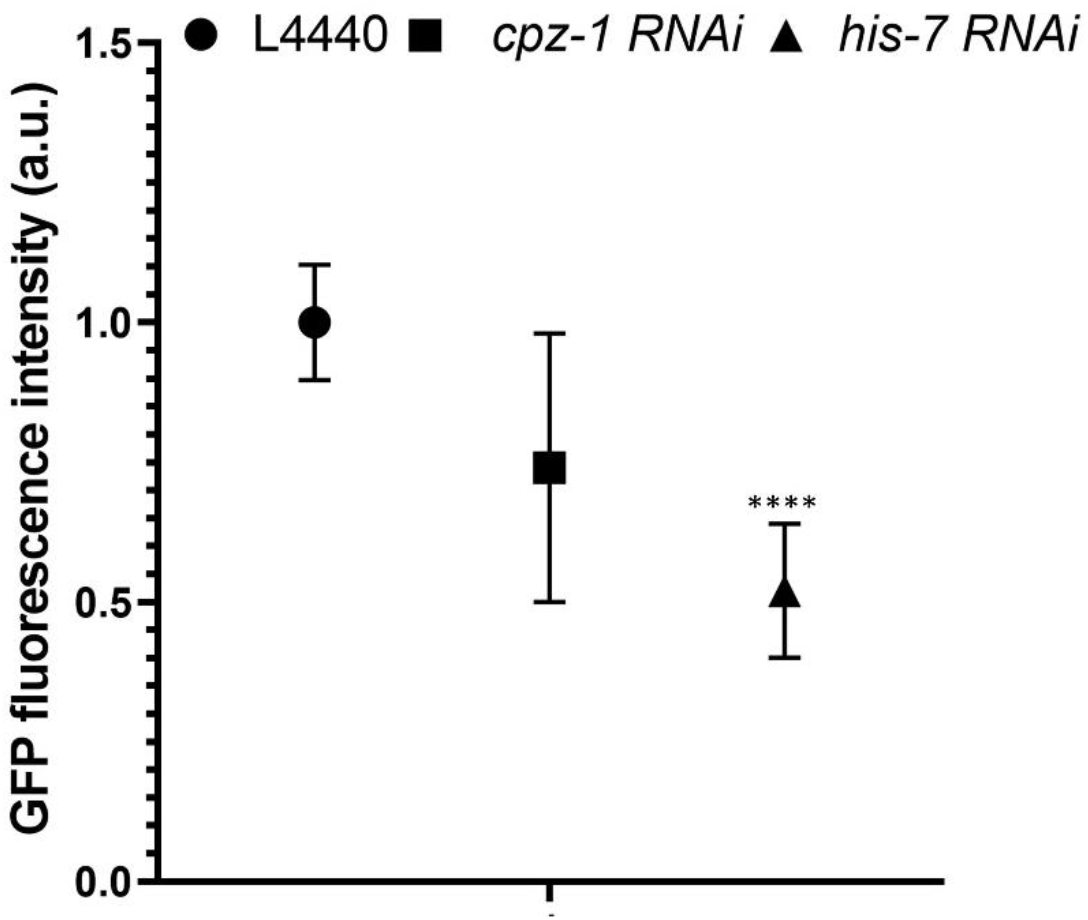
Quantification of fluorescence intensity in *daf-16p::DAF-16::GFP* transgenic animals following RNAi knockdown of *cpz-1*, and *his-7*. Data represents the mean of two replicates (n >15 animals per replicate) and error bars represent the standard deviation. Statistical analyses were done using one-way ANOVA with Dunnett’s post hoc test and significant differences are indicated by stars (*): **** (*p* <0.0001).

### TABLES

**Table S1: List of strains and primers used in the study**.

**Table S2: List of differentially expressed (DE) genes in *pry-1* mutant transcriptome that are associated with reproductive structure development**.

**Table S3: GO-enrichment analysis of 149 DE genes linked to reproductive structure development**.

**Table S4: Tissue and phenotype-enrichment analysis of 149 DE genes linked to reproductive structure development**.

**Table S5: Analysis of DAF-16::GFP localization following RNAi knockdown of *cpz-1* and *his-7* RNAi compared to controls**.

---

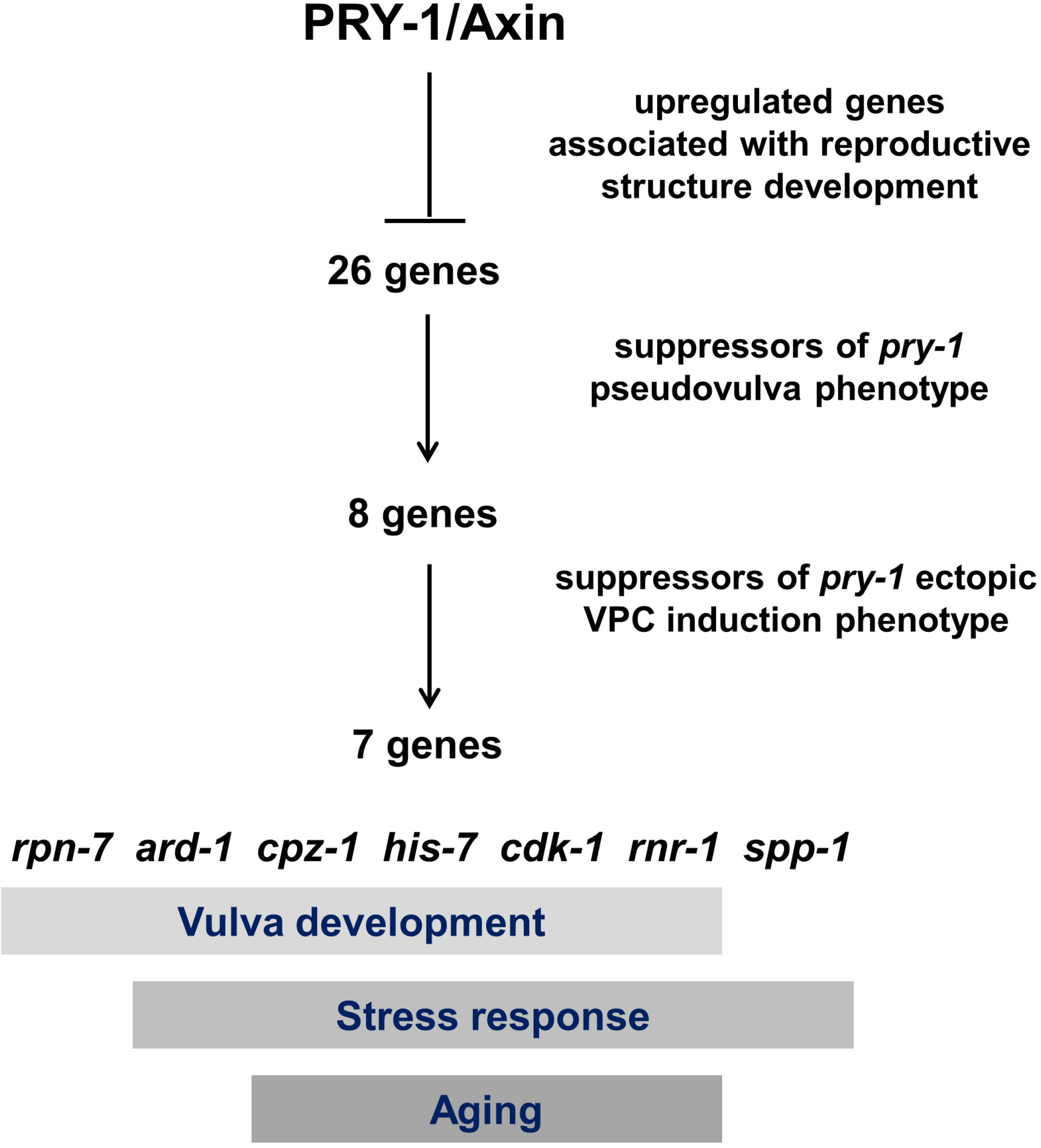

**Aging**
| Gene | Mammalian homolog | GO-biological process | Function in *C. elegans* |
| --- | --- | --- | --- |
| *ard-1* | human HSD17B10 (hydroxysteroid 17-beta dehydrogenase 10) | oxidation-reduction process | Dehydrogenase with NADP binding domain |
| *cdk-1* | Human CDK1 (cyclin dependent kinase 1) | Cell cycle<br>Cell division<br>Protein phosphorylation | protein serine/threonine kinase activity |
| *clsp-1* | human CLSPN (claspin), an adaptor protein that activates ATR kinase | mitotic G2 DNA damage checkpoint<br>activation of protein kinase activity<br>mitotic DNA replication checkpoint | anaphase-promoting complex binding activity; part of the Ataxia telangiectasia and Rad3-related (ATR)/ATL-1 DNA replication checkpoint pathway |
| *cpz-1* | human CTSZ (cathepsin Z) | Vulval development<br>Proteolysis<br>Embryo development ending in birth or egg hatching.<br>Gonad morphogenesis<br>Ecdysis, collagen and cuticulin-based cuticle | Predicted to have cysteine type endopeptidase activity |
| *his-7* | human H2AX (H2A.X variant histone) | DNA repair<br>Chromatin organization<br>Chromatin silencing<br>Regulation of transcription | Predicted to have DNA binding activity |
| *rnr-1* | human RRM1 (ribonucleotide reductase catalytic subunit M1) | DNA replication<br>Metabolic process<br>Deoxyribonucleotide biosynthetic process<br>Oxidation-reduction process | ribonucleoside-diphosphate reductase activity |
| *rpn-7* | human PSMD6 (proteasome 26S subunit, non-ATPase 6) | Proteasome-mediated ubiquitin-dependent protein catabolic process | Component of proteasome 26S complex; ATP-dependent degradation of ubiquitinated proteins |
| *spp-1* | Saposin-B type domain containing protein | Pathogenesis<br>Innate immune response<br>Defense response to Gram-negative/positive bacterium<br>Transmembrane transport | Exhibit channel activity |

